# The magnocellular division of the human medial geniculate nucleus preferentially responds to auditory transients

**DOI:** 10.1101/2022.06.05.494907

**Authors:** Qianli Meng, Keith A. Schneider

## Abstract

The medial geniculate nucleus (MGN), the auditory relay in the thalamus, is composed of three anatomical subdivisions: the ventral, dorsal and medial or magnocellular division. The functional differences among these nuclei have not been studied in humans, and in particular, the function of the magnocellular division is poorly understood in mammals in general. We anatomically segmented the MGN using proton-density-weighted magnetic resonance images (MRI) and measured the functional responses of the MGN to sustained and transient sounds, using functional MRI (fMRI). We observed that voxels in the ventromedial portion of the MGN, corresponding to the magnocellular division, exhibited a strong preference to transient sounds, whereas the remainder of the MGN showed no preference between sustained and transient sounds. We concluded that the magnocellular neurons in the MGN parallel the magnocellular neurons in its visual counterpart, the lateral geniculate nucleus (LGN), and constitute an information stream specialized for encoding stimuli dynamics.

**Significance statement:** The medial geniculate nucleus is the auditory relay in the thalamus. It is composed of three anatomical divisions, of which the function of the magnocellular division is poorly understood. We show, using functional magnetic resonance imaging (fMRI) in humans, that the magnocellular neurons are strongly activated by transient auditory stimuli, similar to the magnocellular neurons in the lateral geniculate nucleus, the visual thalamic relay, which are rapidly adapting and specialized to encode visual transients. These results confirm that the auditory system represents stimuli using parallel information streams, employing similar encoding strategies as in other sensory modalities.

## Introduction

The medial geniculate nucleus (MGN), the auditory relay nucleus in the thalamus, has three main subdivisions based on cellular morphology: the ventral, dorsal and medial divisions (Jones, 2003). The latter is hereafter referred to as the magnocellular division based on a population of large neuron cell bodies found here (Winer and Morest, 1983; Winer, 1984), comparable to the magnocellular neurons in the visual lateral geniculate nucleus (LGN). The magnocellular and ventral divisions of the MGN contain tonotopic maps, orderly representations of sound frequency (Jones, 2003; Moerel et al., 2015; Glad Mihai et al., 2019). Little is otherwise known about the response properties of neurons in the magnocellular division of the MGN in humans; the neurons here in other species habituate rapidly (Aitkin, 1973; Bäuerle et al., 2011) and exhibit short response latencies (Anderson and Linden, 2011).

The LGN in comparison has been thoroughly studied, in humans and particularly in other primates. In the macaque, magnocellular neurons in the LGN are specialized to encode transient visual stimuli (Derrington and Lennie, 1984; Maunsell et al., 1999; Solomon et al., 2004). It seems to be the case that magnocellular neurons throughout the sensory and motor systems of the brain are specialized for temporal processing (Stein and Walsh, 1997; Trussell, 1997; Stein and Talcott, 1999; Stein, 2001). We therefore hypothesized that magnocellular neurons in the MGN might be analogous to those in the LGN and would selectively encode auditory transients.

## Materials and Methods

### Subjects

11 subjects (nine female) participated, all 18–32 years old. None had any neurological disorders, and all were right-handed. The subjects provided informed written consent under the research protocol approved by the Institutional Review Board at the University of Delaware.

### Stimuli

The stimuli were generated using MATLAB software (The MathWorks, Inc.) with the Psychophysics Toolbox 3 functions (Brainard, 1997; Pelli, 1997; Kleiner et al., 2007) running on a Linux computer. The stimuli were synchronized to the MRI acquisition using a trigger signal from the scanner, interfaced to the computer through a fORPs response box (Current Designs, Inc.). Auditory stimuli were presented through headphones (OptoActive II, Optoacoustics Ltd.) with real-time algorithmic, out-of-phase harmonic active noise cancelation that attenuated the background noise from the scanner. All subjects passively listened to the stimuli and reported clearly hearing them throughout the scanning procedures. Twelve blocks each of sustained or transient auditory stimuli were presented, randomly interleaved during each 312 s scanning run (Fig. 1). Blocks were 13 s in duration, consisting of approximately 9 s of sound and the remaining approximately 4 s of the block unstimulated (silent). The stimuli in each block were sampled from broadband natural sounds with no linguistic content, e.g. bells, wild animal roars, car horns, trumpets and steam engine whistles. The transient blocks consisted of a series of three to four sound bursts of the samples, separated by 0.5 s of silence and windowed with a square wave to produce abrupt onsets and offsets (Fig. 1). The duration of these bursts was either 50– 150 ms for shorter bursts or 200–300 ms for longer bursts. The silent gap between bursts was 20–40 ms. This timing was chosen to approximate the typical syllabic timing in speech. The sustained blocks were composed from the same sound samples, but 0.5–2 s in duration, separated by 0.5 s and windowed with sine onset and offset ramps, each one-quarter of the sound duration, to eliminate transients. Visual stimuli were also presented, consisting of high contrast checkerboards flickering at various frequencies, with timing independent of the auditory stimuli, but these were not analyzed for the present study.

**Figure. 1.**
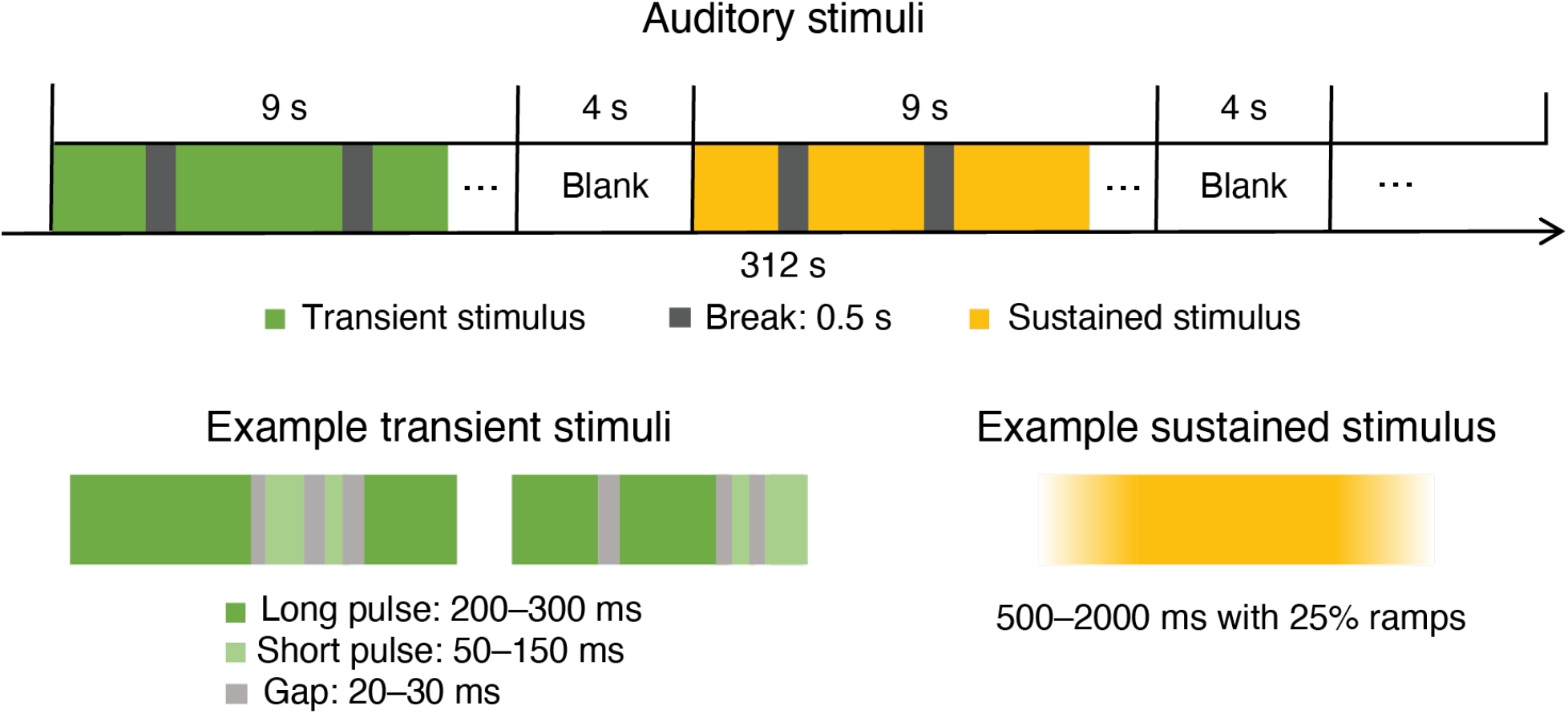
Experimental design and mean activations. Transient and sustained auditory stimuli were randomly interleaved in separate 13 s blocks that included approximately 9 s of sound and 4 s of silence. The transient stimuli consisted of short (50–150 ms) or long (200–300 ms) pulses with a 20–40 ms gap between. The sustained stimuli were 500– 2000 ms in duration with 125– 500 ms sine ramped onsets and offsets.

### Neuroimaging

Data were acquired using a 3T Siemens (Erlangen, Germany) Prisma MRI scanner with a 64-channel head coil at the Center for Biomedical and Brain Imaging at the University of Delaware. All subjects participated in two to five MRI scanning sessions. A high-resolution T_1_-weighted scan was acquired for each subject in each scanning session [MPRAGE, TR = 2080 ms, TE = 4.64 ms, flip angle = 9°, 208 sagittal slices, FOV = 210 × 210 mm, acquisition matrix = 288 × 288, isotropic (0.7 mm)^3^ resolution, parallel imaging (iPAT) acceleration factor (GRAPPA) = 2]. In one session, we acquired 40 proton density-weighted (PD) spin-echo structural images [acquisition time 89 s, TR = 3000 ms, TE = 16 ms, flip angle = 150°, 35 coronal 1 mm thick slices covering the thalamus, FOV = 256 × 256 mm, acquisition matrix = 256 × 256, isotropic 1 mm^3^ resolution, iPAT GRAPPA = 2]. In the remaining sessions, we acquired six to ten runs, 209 volumes each, of functional data using a multi-band gradient echo EPI sequence [TR = 1.5 s, TE = 39 ms, flip angle = 45°, 84 horizontal 1.5 mm thick slices, FOV = 192 × 192 mm, acquisition matrix = 128 × 128, isotropic (1.5 mm)^3^ resolution, A → P phase encoding, partial Fourier factor = 6/8, slice acceleration factor = 6, bandwidth = 1562 Hz/Px]. The subjects’ heads were surrounded by foam padding to reduce head movements.

### Anatomical regions of interest

The location of the MGN in humans is well known from functional and anatomical studies (Yetkin et al., 2004; Devlin et al., 2006; Jiang et al., 2013; García-Gomar et al., 2019; Sitek et al., 2019). The 40 PD images were registered using an affine transformation to correct for displacement between acquisitions, up-sampled to twice the resolution in each dimension and averaged to create a mean image with high signal-to-noise. These images were aligned to the T_1_ and used to manually trace the anatomical extent of each MGN (Fig. 2). These anatomical ROIs were used to guide the functional ROI, and the two ROIs matched well except in one subject (Fig. 2).

**Figure 2.**
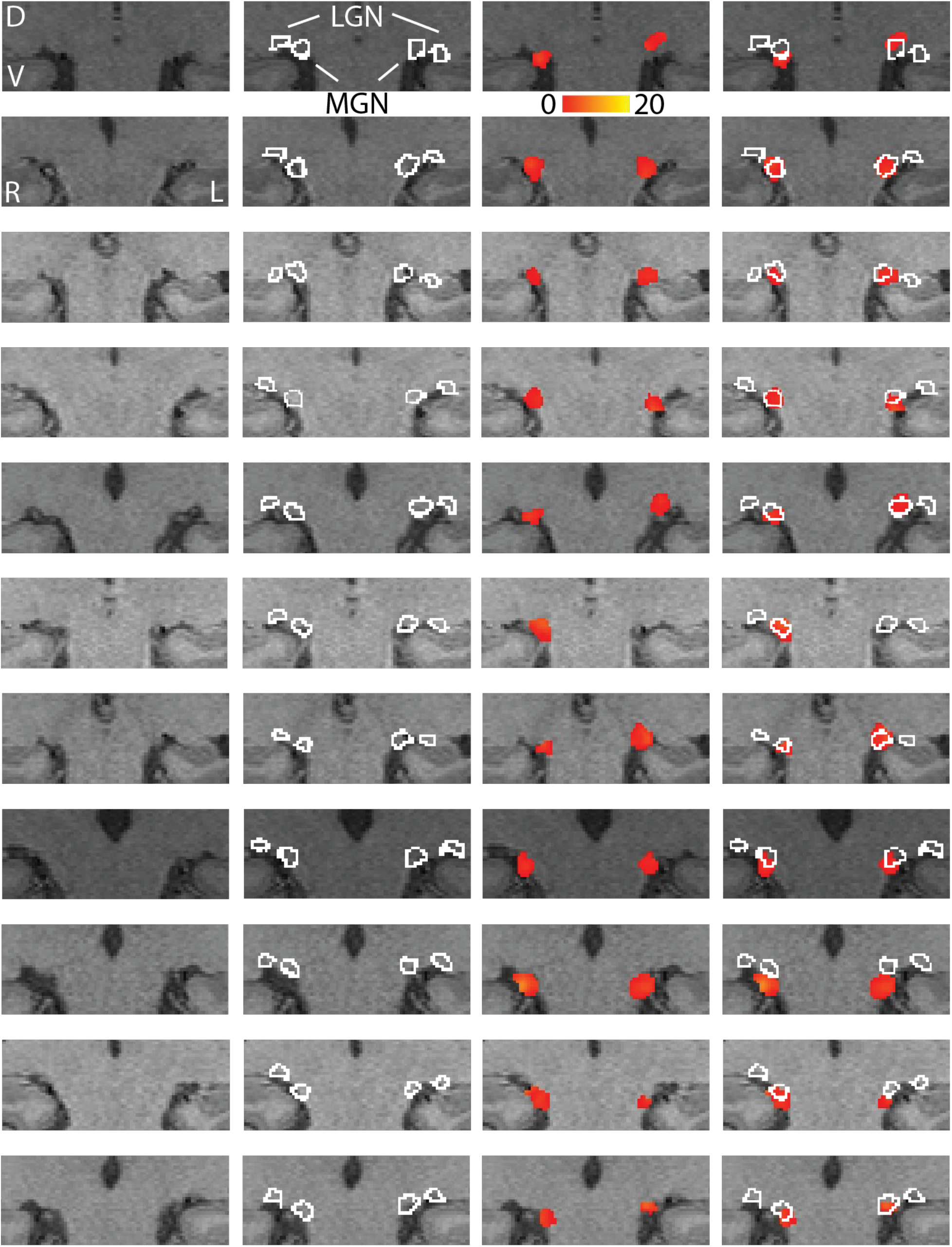
Region of interest selection. Each subject is shown as a separate row. For each subject, a zoomed coronal slice is shown through the thalamus, with the subject’s structural image as the background. The columns from left to right show: 1. structural image, 2., outlines of the anatomical segmentations of the MGN and LGN, based on proton-density images, 3. functional activation to combined auditory stimuli, with the color bar indicating for each voxel the z-score of the sound vs. blank contrast, 4. overlay of the anatomical ROI on the functional activations. D, dorsal; V, ventral; R, right; L, left.

### Experimental design and statistical analysis

Individual subjects were analyzed in their native space. Functional data were preprocessed using FEAT (FMRI Expert Analysis Tool) Version 6.00, part of FSL (FMRIB’s Software Library, www.fmrib.ox.ac.uk/fsl)(Jenkinson et al., 2012). The functional images were realigned to correct for small head movements using FLIRT (Jenkinson et al., 2002) and then linearly registered to each participant’s T_1_. Each image was smoothed with a 2 mm full width at half maximum (FWHM) Gaussian kernel. Each subject’s data were analyzed with a general linear model (Woolrich et al., 2001), with two explanatory variables (EVs) accounting for the transient and sustained stimuli, with the silent periods as the baseline. The estimated motion parameters for each run were included as covariates of no interest, and the motion outliers were defined as additional confound EVs. A fixed effects analysis combined the multiple scanning runs within each subject. Because the amount of data varied among subjects (who had one to four functional scanning sessions each), the noise level among subjects also varied. Therefore, to define the functional ROI, a cluster-based significance test was used with a different height threshold for each subject, with 𝓏∈ {0.5, 1, 1.5, 2} (the lowest necessary for the noisiest three of the 22 MGN; one other MGN was not detected), and a cluster *p* < .05, corrected for multiple comparisons (Worsley et al., 2002). This resulted in a functional activation distinct from the background noise and matching the size of the anatomical ROI. A transient index, (*T* − *S*)/ (*T* + *S*), was computed for each activated voxel using the weights of the transient (*T*) and sustained (*S*) activations. To compare the activation in the whole MGN across subjects, the mean activations for each EV were calculated over all activated voxels in each MGN, and these mean activations were subjected to a mixed effects repeated measures model with hemisphere (left or right) and stimulus (S or T) as within-subjects fixed effects.

The group analysis was computed in standard space in the same manner as the individual analyses, with the following differences. Registration from the T_1_ to standard space was calculated using FNIRT nonlinear registration (Andersson et al., 2007). A mixed effects (FLAME 1+2) analysis was conducted to compute the group EVs from each subject’s individual results (Woolrich et al., 2004). The group transient index was computed for each voxel using the weights of the group EVs. The ROIs of the MGN in standard space were defined using the Jülich histological atlas (Bürgel et al., 2006).

## Results

The MGN was segmented in each subject using structural imaging, and the mean volume across subjects was 134.2 ± 7.0 mm^3^ (mean ± SEM). Using functional magnetic resonance imaging (fMRI), we measured the relative responses to transient and sustained auditory stimuli within the MGN. We imaged subjects as they passively listened to transient or sustained non-linguistic sounds, using a block design (Fig. 1). The MGN in all subjects were well activated by the combined auditory stimuli (Fig. 2), bilaterally in all but one subject. In the whole MGN, i.e., the mean of the activated voxels in each MGN in the native spaces of each subject, we found that subjects exhibited greater activation to the transient than sustained stimuli (*F*_1,7.8_ = 259.3, *p* < .0001, Fig. 3B), with no difference between the left and right MGN (*F*_1,9.9_ = 0.26, *p* = .6202).

**Figure 3.**
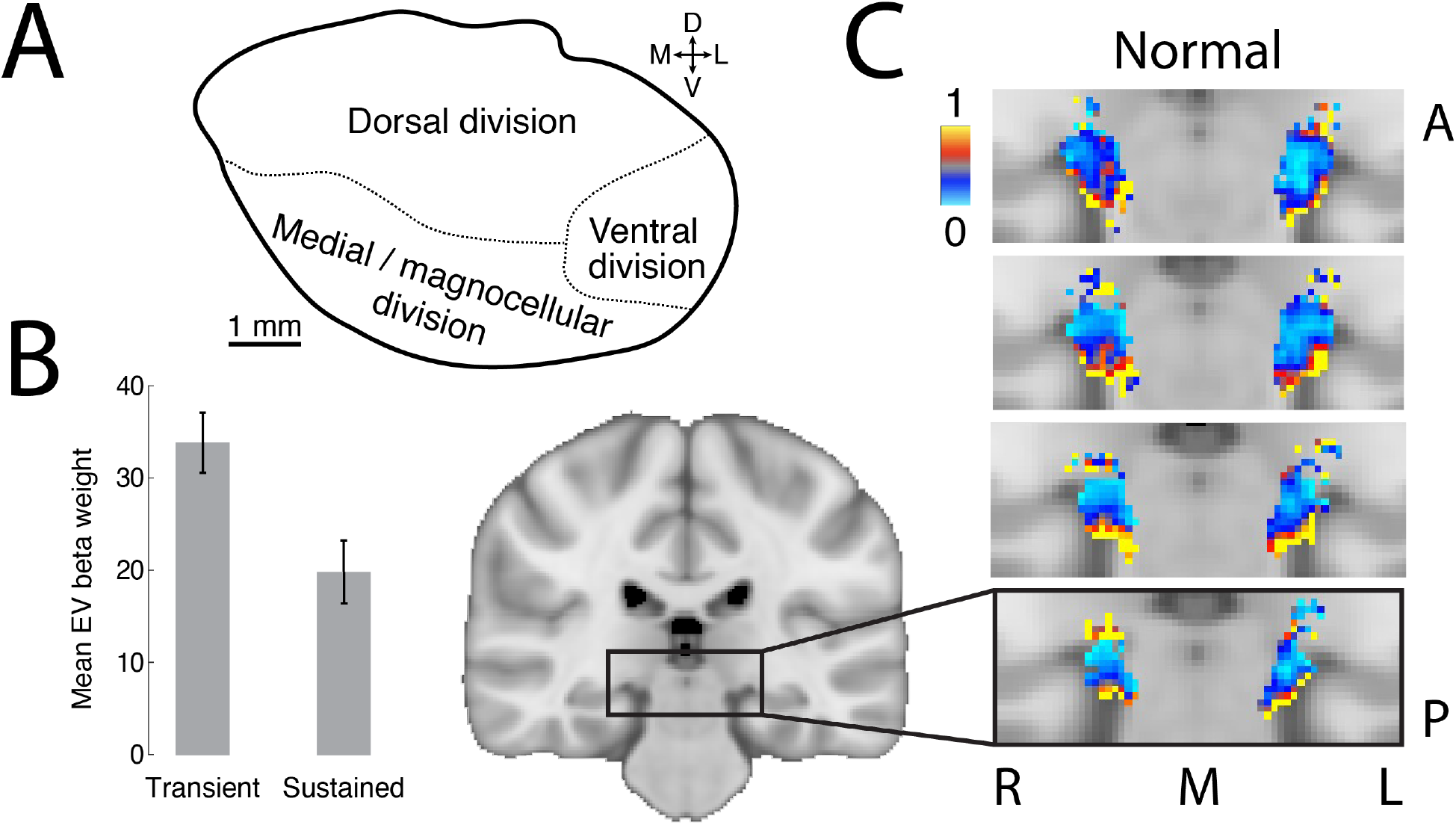
Group results. A. A schematic of the divisions of the human MGN, based on cellular morphology (Winer, 1984). The medial division is also known as the magnocellular division (D, dorsal; V, ventral; M, medial; L lateral). B. The mean transient (*T*) and sustained (*S*) activations across subjects are shown, collapsed across left and right MGN in the native space of each subject. C. A zoomed region of the brain is displayed covering the thalamus on four adjacent coronal slices, arranged anterior (A) to posterior (P). The transient index, (*T* − *S*)/ (*T* + *S*), is displayed for each voxel in the MGN in standard space, computed over all subjects, where *T* and *S* are the transient and sustained group activations, respectively. The blue voxels with indices near zero showed no preference between the transient and sustained stimuli. The red and yellow voxels with indices closer to one were more strongly activated by the transient stimuli. Voxels with large transient indices were largely confined to the ventromedial portion of the MGN, the magnocellular division (L, left; R, right; M, midline).

To visualize the distribution of activations to the transient (*T*) and sustained (*S*) stimuli within the MGN volume, we computed a normalized transient index (*T* − *S*)/ (*T* + *S*) for each voxel. In both the group in standard space (Fig. 3C) and in every subject individually in their native spaces (Fig. 4), we observed strong responses to the transient stimuli (high transient indices) along the entire ventral surface of the MGN, corresponding to the expected size and location of the magnocellular division as expected from the anatomy (Fig. 3A). There were also small clusters of increased activation to transient vs. sustained stimuli on the dorsolateral surface of the MGN. There were no areas in the cortex selective for the transient stimuli (Fig. 5).

**Figure 4.**
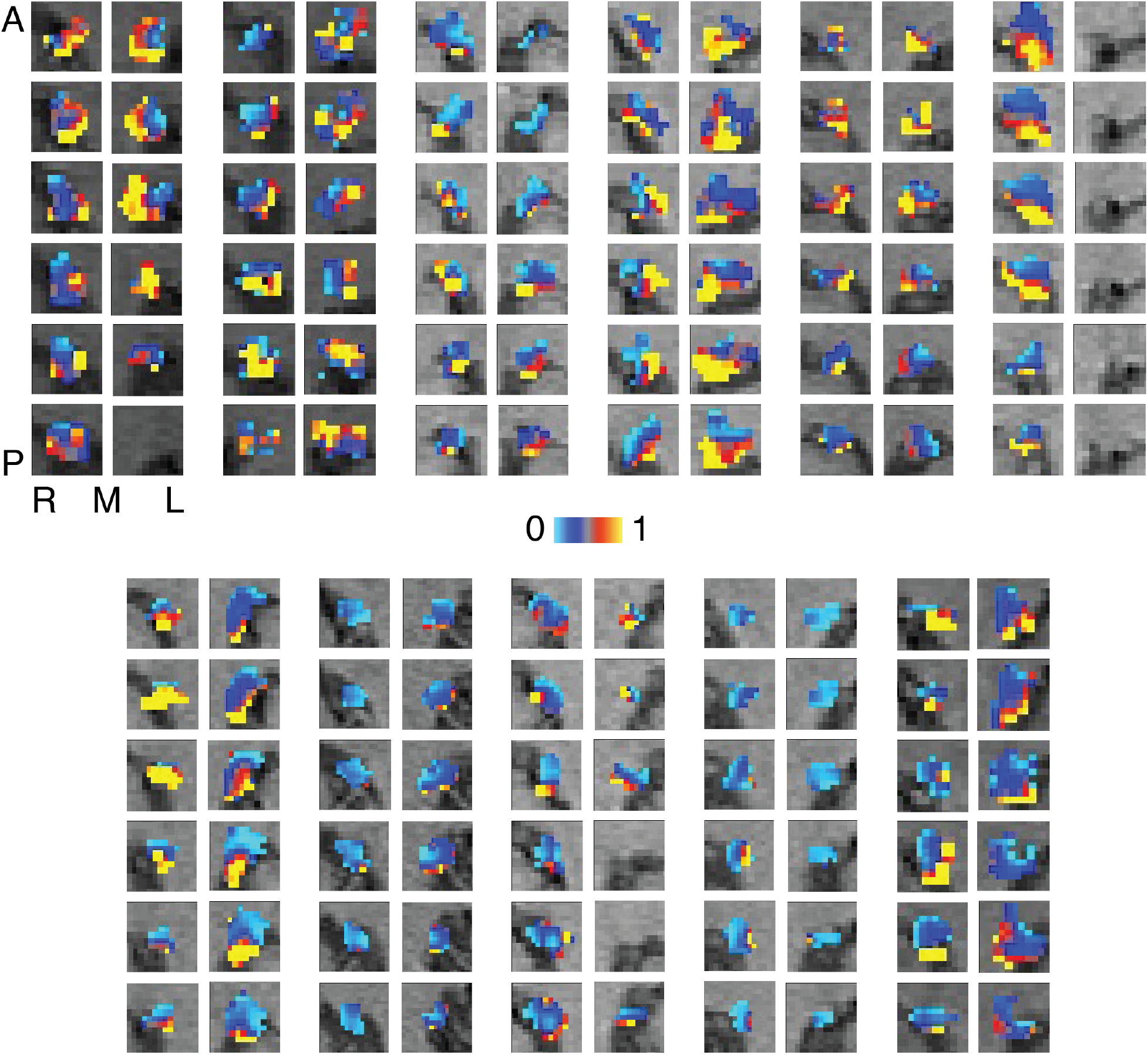
Transient indices for all individual subjects in native space. Each pair of columns shows the right (R) and left (L) MGN for a single subject, arranged relative to the midline (M), with six coronal slices per MGN, ordered anterior (A) to posterior (P). The transient index is displayed for each activated voxel in the MGN in each subject. The blue voxels have a transient index near zero, indicating that there was little or no difference in response to the transient or sustained stimuli, whereas red and yellow voxels exhibited a stronger response to the transient stimuli.

**Figure 5.**
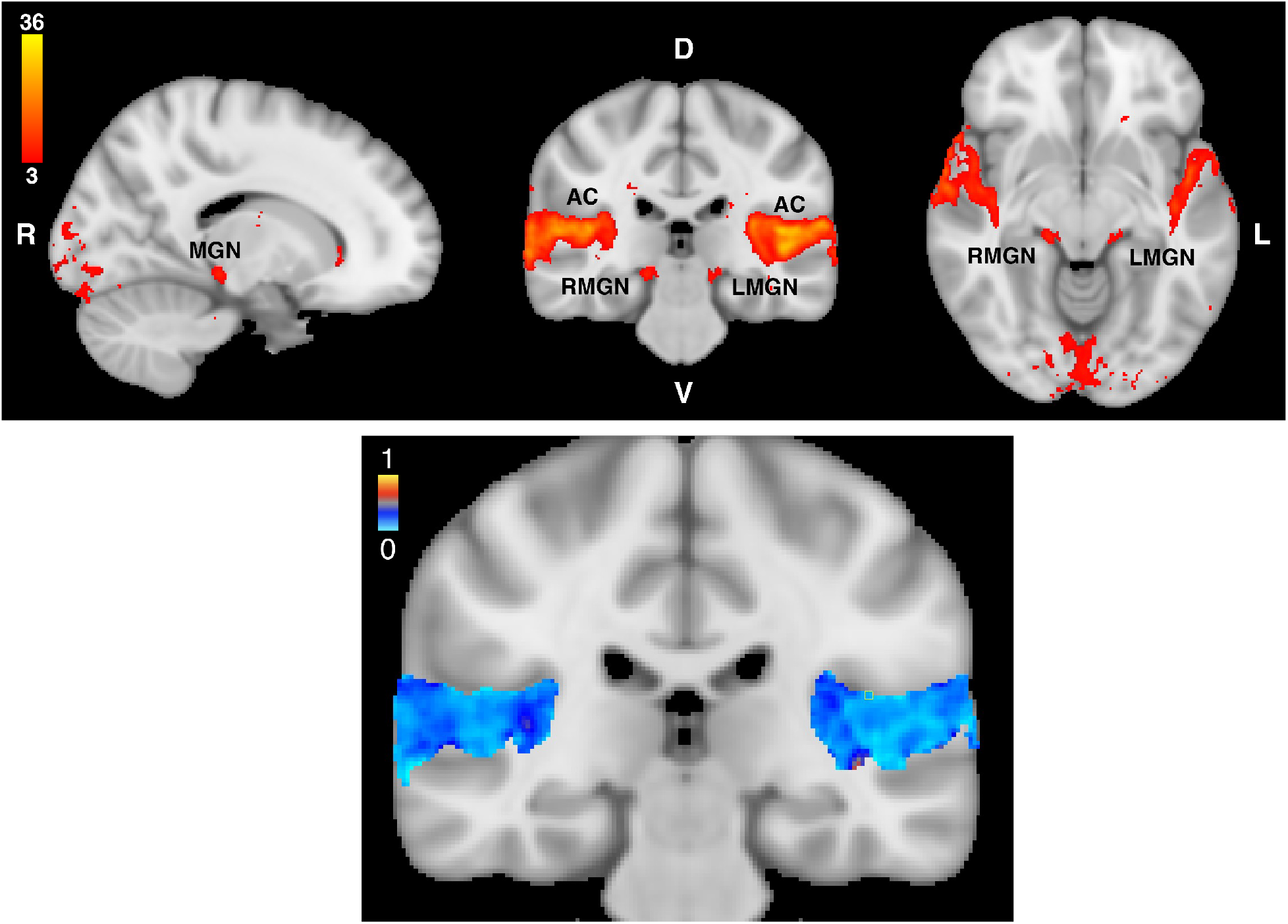
Cortical activations. In the top panel, group activations to the combined auditory stimuli are shown. *z* > 3, *p* < .05. D, dorsal; V, ventral; R, right; L, left; AC, auditory cortex; MGN, medial geniculate nucleus. In the bottom panel, the group transient indices in the auditory cortex are shown.

## Discussion

Across subjects, we observed a strong and remarkably consistent preference for transient compared to sustained auditory stimuli in the ventromedial portion of the MGN, corresponding to its magnocellular division. The remainder of the MGN responded equally to sustained or transient stimuli. Although the whole MGN can be readily segmented using anatomical imaging, and the location of the magnocellular division is generally known, we show that a contrast between transient and sustained stimuli can be used as an effective functional localizer for the magnocellular division.

Throughout the sensory and motor systems in the brain, there are neurons with large cell bodies that are specialized for temporal processing (Stein and Walsh, 1997; Stein and Talcott, 1999; Stein, 2001), comprising a magnocellular “system”. Although the magnocellular division in the human MGN contains a variety of different neuron types (Winer, 1984), the functional response seems to be dominated by the magnocellular neurons, as found in the magnocellular divisions in the other auditory relay nuclei (Trussell, 1997).

The function of the magnocellular portion of the MGN is interesting to consider. In the visual system, the magno- and parvocellular divisions of the LGN comprise parallel streams that are optimized to encode different aspects of the visual stimuli. The visual system is thought to diverge into parallel channels that specialize in localization or recognition (Goodale and Milner, 1992). The auditory system is not thought to be organized in this way, but two channels encoding transient and sustained portions of an auditory signal might be optimal to reduce overall activation yet still permit precise encoding of rapidly modulating signals, such as speech, that convey important information.

## Acknowledgements

We would like to acknowledge funding from National Institutes of Health (National Eye Institute) grant 1R01EY028266 (KAS), Joy Lin and Julia Kausel for their assistance with the MRI scanning, and Anton Lebed for his assistance in analyzing the data.

## References

Aitkin LM (1973) Medial geniculate body of the cat: responses to tonal stimuli of neurons in medial division. J Neurophysiol 36:275–283 Available at: https://www.physiology.org/doi/10.1152/jn.1973.36.2.275.

Anderson LA, Linden JF (2011) Physiological differences between histologically defined subdivisions in the mouse auditory thalamus. Hear Res 274:48–60 Available at: http://dx.doi.org/10.1016/j.heares.2010.12.016.

Andersson JLR, Jenkinson M, Smith S (2007) Non-linear registration, aka spatial normalization (FMRIB technical report TR07JA2).

Bäuerle P, von der Behrens W, Kössl M, Gaese BH (2011) Stimulus-Specific Adaptation in the Gerbil Primary Auditory Thalamus Is the Result of a Fast Frequency-Specific Habituation and Is Regulated by the Corticofugal System. J Neurosci 31:9708–9722 Available at: https://www-jneurosci-org.udel.idm.oclc.org/content/31/26/9708 [Accessed May 9, 2022].

Brainard DH (1997) The Psychophysics Toolbox. Spat Vis 10:433–436 Available at: http://www.ncbi.nlm.nih.gov/entrez/query.fcgi?cmd=Retrieve&db=PubMed&dopt=Citation&list_uids=9176952.

Bürgel U, Amunts K, Hoemke L, Mohlberg H, Gilsbach JM, Zilles K (2006) White matter fiber tracts of the human brain: Three-dimensional mapping at microscopic resolution, topography and intersubject variability. Neuroimage 29:1092–1105.

Derrington AM, Lennie P (1984) Spatial and temporal contrast sensitivities of neurones in lateral geniculate nucleus of macaque. J Physiol 357:219–40 Available at: http://www.ncbi.nlm.nih.gov/entrez/query.fcgi?cmd=Retrieve&db=PubMed&dopt=Citation&list_uids=6512690.

Devlin JT, Sillery EL, Hall DA, Hobden P, Behrens TEJ, Nunes RG, Clare S, Matthews PM, Moore DR, Johansen-Berg H (2006) Reliable identification of the auditory thalamus using multi-modal structural analyses. Neuroimage 30:1112–20 Available at: http://www.ncbi.nlm.nih.gov/entrez/query.fcgi?cmd=Retrieve&db=PubMed&dopt=Citation&list_uids=16473021.

García-Gomar MG, Strong C, Toschi N, Singh K, Rosen BR, Wald LL, Bianciardi M (2019) In vivo probabilistic structural atlas of the inferior and superior colliculi, medial and lateral geniculate nuclei and superior olivary complex in humans based on 7 tesla MRI. Front Genet 10.

Glad Mihai P, Moerel M, De Martino F, Trampel R, Kiebel S, Von Kriegstein K (2019) Modulation of tonotopic ventral medial geniculate body is behaviorally relevant for speech recognition. Elife 8.

Goodale MA, Milner AD (1992) Separate visual pathways for perception and action. Trends Neurosci 15:20–25 Available at: http://www.ncbi.nlm.nih.gov/entrez/query.fcgi?cmd=Retrieve&db=PubMed&dopt=Citation&list_uids=1374953.

Jenkinson M, Bannister P, Brady M, Smith S (2002) Improved optimisation for the robust and accurate linear registration and motion correction of brain images. Neuroimage 17:825–841.

Jenkinson M, Beckmann CF, Behrens TEJ, Woolrich MW, Smith SM (2012) FSL. Neuroimage 62:782–790 Available at: https://linkinghub.elsevier.com/retrieve/pii/S1053811911010603.

Jiang F, Stecker GC, Fine I (2013) Functional localization of the auditory thalamus in individual human subjects. Neuroimage 78:295–304 Available at: http://dx.doi.org/10.1016/j.neuroimage.2013.04.035.

Jones EG (2003) Chemically Defined Parallel Pathways in the Monkey Auditory System. Ann N Y Acad Sci 999:218–233 Available at: https://onlinelibrary-wiley-com.udel.idm.oclc.org/doi/full/10.1196/annals.1284.033 [Accessed May 9, 2022].

Kleiner M, Brainard D, Pelli D, Ingling A, Murray R, Broussard C (2007) Perception. [Pion Ltd.]. Available at: https://nyuscholars.nyu.edu/en/publications/whats-new-in-psychtoolbox-3 [Accessed September 12, 2018].

Maunsell JHR, Ghose GM, Assad JA, McAdams CJ, Boudreau CE, Noerager BD (1999) Visual response latencies of magnocellular and parvocellular LGN neurons in macaque monkeys. Vis Neurosci 16:1–14 Available at: http://www.ncbi.nlm.nih.gov/pubmed/10022474 [Accessed February 28, 2019].

Moerel M, De Martino F, Uğurbil K, Yacoub E, Formisano E (2015) Processing of frequency and location in human subcortical auditory structures. Sci Rep 5:17048 Available at: http://www.nature.com/scientificreports [Accessed July 13, 2020].

Pelli DG (1997) The VideoToolbox software for visual psychophysics: transforming numbers into movies. Spat Vis 10:437–442 Available at: https://brill.com/view/journals/sv/10/4/article-p437_16.xml.

Sitek KR, Gulban OF, Calabrese E, Johnson GA, Lage-Castellanos A, Moerel M, Ghosh SS, De Martino F (2019) Mapping the human subcortical auditory system using histology, postmortem MRI and in vivo MRI at 7T. Elife 8 Available at: https://elifesciences.org/articles/48932 [Accessed July 13, 2020].

Solomon SG, Peirce JW, Dhruv NT, Lennie P (2004) Profound Contrast Adaptation Early in the Visual Pathway. Neuron 42:155–162 Available at: http://www.ncbi.nlm.nih.gov/entrez/query.fcgi?cmd=Retrieve&db=PubMed&dopt=Citation&list_uids=15066272.

Stein J (2001) The magnocellular theory of developmental dyslexia. Dyslexia 7:12–36 Available at: http://www.ncbi.nlm.nih.gov/entrez/query.fcgi?cmd=Retrieve&db=PubMed&dopt=Citation&list_uids=11305228.

Stein J, Talcott J (1999) Impaired neuronal timing in developmental dyslexia—the magnocellular hypothesis. Dyslexia 5:59–77 Available at: https://onlinelibrary.wiley.com/doi/10.1002/(SICI)1099-0909(199906)5:2%3C59::AID-DYS134%3E3.0.CO;2-F [Accessed May 9, 2022].

Stein J, Walsh V (1997) To see but not to read; the magnocellular theory of dyslexia. Trends Neurosci 20:147–52 Available at: http://www.ncbi.nlm.nih.gov/entrez/query.fcgi?cmd=Retrieve&db=PubMed&dopt=Citation&list_uids=9106353.

Trussell LO (1997) Cellular mechanisms for preservation of timing in central auditory pathways. Curr Opin Neurobiol 7:487–492 Available at: https://linkinghub.elsevier.com/retrieve/pii/S095943889780027X [Accessed May 9, 2022].

Winer JA (1984) The human medial geniculate body. Hear Res 15:225–247 Available at: https://pubmed.ncbi.nlm.nih.gov/6501112/ [Accessed July 13, 2020].

Winer JA, Morest DK (1983) The medial division of the medial geniculate body of the cat: Implications for thalamic organization. J Neurosci 3:2629–2651.

Woolrich MW, Behrens TEJ, Beckmann CF, Jenkinson M, Smith SM (2004) Multilevel linear modelling for FMRI group analysis using Bayesian inference. Neuroimage 21:1732–1747.

Woolrich MW, Ripley BD, Brady M, Smith SM (2001) Temporal autocorrelation in univariate linear modeling of FMRI data. Neuroimage 14:1370–1386.

Worsley KJ, Liao CH, Aston J, Petre V, Duncan GH, Morales F, Evans AC (2002) A general statistical analysis for fMRI data. Neuroimage 15:1–15 Available at: http://www.ncbi.nlm.nih.gov/pubmed/11771969.

Yetkin FZ, Roland PS, Mendelsohn DB, Purdy PD (2004) Functional magnetic resonance imaging of activation in subcortical auditory pathway. Laryngoscope 114:96–101 Available at: http://www.ncbi.nlm.nih.gov/entrez/query.fcgi?cmd=Retrieve&db=PubMed&dopt=Citation&list_uids=14710002.

